# Virtual Immersion in Biomedical Engineering (VIBE): Exposing undergraduates to culturally sensitive engineering design and professional experiences at scale

**DOI:** 10.64898/2026.02.08.704621

**Authors:** Kristen Billiar, Taimoor Afzal, Olufunmilayo Ayobami, Solomon Mensah

## Abstract

**Challenge:** Undergraduate engineering students often face two major barriers: limited access to culturally sensitive global experiences and difficulty securing design internship opportunities. Traditional study abroad and internship programs can be inaccessible due to financial, geographic, or structural constraints, hindering the development of global competencies and practical engineering skills.

**Novel Initiative:** To address these gaps, we created the Virtual Immersion in Biomedical Engineering (VIBE) program, a free, inclusive, and scalable summer experience. The 8-week online program blends interactive sessions on the engineering design process (EDP), faculty led research talks, and professional development workshops. Collaborative team-based design challenges focused on global health and healthcare disparities are completed by international teams of 3-4 students. From 2021 to 2024, more than 800 students from over 20 countries participated, with an even balance of male and female students. Student presentations (four-minute videos uploaded to YouTube) were evaluated using rubrics with the aid of a large language model. The assessment revealed substantial learning of the EDP, especially in problem definition and communication.

**Reflection:** Participant feedback underscored the program’s benefits: exposure to global collaboration, professional development, and flexibility. However, common challenges included time zone coordination and team dynamics. Despite these hurdles, VIBE has demonstrated that virtual experiential learning can effectively build engineering design skills and global awareness. It offers a promising, accessible alternative to traditional high-cost global programs, expanding equitable opportunities for engineering students worldwide.

## Challenge Statement

We addressed two related challenges in this work, both preferentially affecting minoritized groups in the United States (U.S.) and students from low- and middle-income countries (LMICs), and both exacerbated by the COVID-19 pandemic. First, the difficulty for engineering students in obtaining a global perspective and cultural sensitivity, and second, the limited availability of summer engineering design experiences and opportunities for professional development.

Global perspectives and authentic design experiences that enhance cultural competence are crucial for engineering students’ success in an increasingly interconnected world. Yet international experiences, such as semester- or year-long study abroad programs, are difficult to fit into packed engineering curricula without disrupting time to graduation, leading to less than 4% of engineering students participating in semesters abroad (Renganathan, Gerhardt et al. 2008, Yates, Wentz et al. 2020). International programs are especially difficult to access for U.S. students from populations without sufficient funds or knowledge of opportunities (Harris and Hynes 2019). Students from other countries also face significant barriers to obtaining international engineering experiences, primarily due to financial, bureaucratic, and institutional constraints(Keshavamurthy, Minerick et al. 2008, Kjellgren and Richter 2022).

Summer internships in major-related industries and research laboratories are more sought after than international experiences but are difficult to get. Further first-year, first-generation, and low-income students access internships at lower rates compared to their peers (Yang, Towles et al. 2024). Additionally, many summer programs that are available do not offer professional development opportunities or broader perspectives about the biomedical engineering field.

### Novel Initiative

To meet these two related challenges, we created the Virtual Immersion in Biomedical Engineering (VIBE) an 8-week summer program which provides authentic bioengineering design experiences for international teams of students in a virtual environment freely accessible to students from all institutions. To provide a broader perspective, the program also includes weekly research presentations and conversations with BME faculty, and professional development workshops.

Our approach to learning is rooted in constructivist theory which requires learners to actively co-construct meaning with peers (McHenry, Depew et al. 2005). This collaborative sense-making process requires learners to extend themselves beyond simple memorization and to use higher-order thinking skills, such as the applying, analyzing, and creating levels of Bloom’s taxonomy revised (Bloom, Engelhart et al.1956, Anderson, Krathwohl et al. 2001). For example, through open-ended problem solving, VIBE participants applied knowledge from lectures on the engineering design process (EDP) to their own team-based virtual design projects. The authentic nature of the problems and the element of choice in approaches and solutions are key aspects of the project-based learning (PjBL) pedagogical framework that add to the benefits of problem-based learning (PBL) (Bell 2010, Clyne and Billiar 2016).

To recruit a diverse audience of students from across the United States and internationally (with a focus on underserved populations), we promote the program on a webpage, [redacted] advertise the program through email listserves (e.g., the BME/BioE Council of Chairs), and sent emails to our colleagues’ international contacts in Ghana, India, Pakistan, and Nigeria. In addition to these channels, the program is advertised on the department of biomedical engineering faculty’s LinkedIn profiles. After the first year in which over 300 students participated, about 10% of the applicants in the following year learned of the program from their peers from the previous cohort. The demographics of the participants over the past four years (2021-2024) are provided in ***Table I***.

**Table I:**
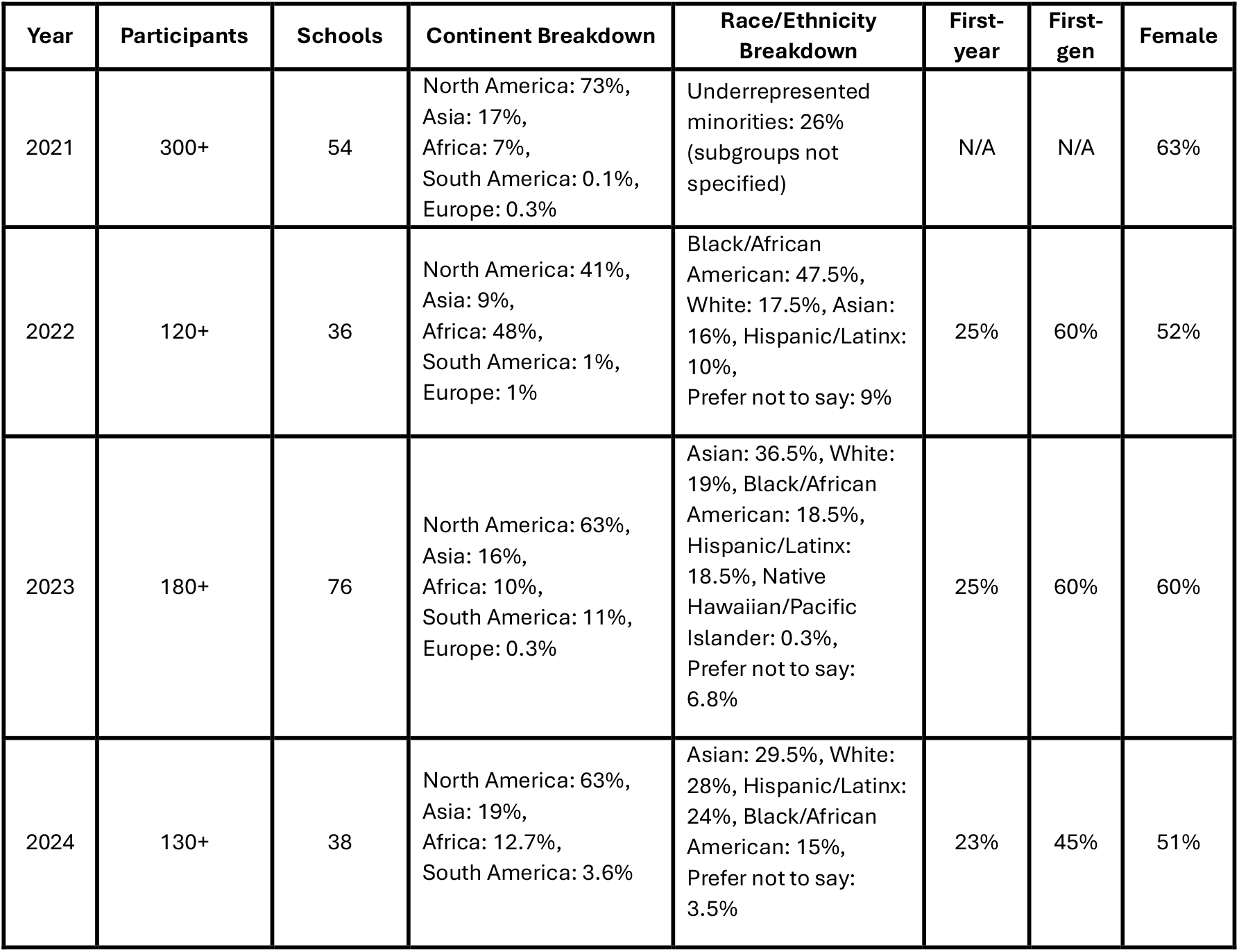
Numbers of participants who successfully completed the VIBE program each year and the number of schools (universities/colleges) and continents where they are located, their demographics, and the proportion of first year, first-generation, and female students.

| Year | Participants | Schools | Continent Breakdown | Race/Ethnicity Breakdown | First-year | First-gen | Female |
| --- | --- | --- | --- | --- | --- | --- | --- |
| 2021 | 300+ | 54 | North America: 73%,<br>Asia: 17%,<br>Africa: 7%,<br>South America: 0.1%,<br>Europe: 0.3% | Underrepresented minorities: 26% (subgroups not specified) | N/A | N/A | 63% |
| 2022 | 120+ | 36 | North America: 41%,<br>Asia: 9%,<br>Africa: 48%,<br>South America: 1%,<br>Europe: 1% | Black/African American: 47.5%,<br>White: 17.5%, Asian: 16%,<br>Hispanic/Latinx: 10%,<br>Prefer not to say: 9% | 25% | 60% | 52% |
| 2023 | 180+ | 76 | North America: 63%,<br>Asia: 16%,<br>Africa: 10%,<br>South America: 11%,<br>Europe: 0.3% | Asian: 36.5%, White: 19%,<br>Black/African American: 18.5%,<br>Hispanic/Latinx: 18.5%,<br>Native Hawaiian/Pacific Islander: 0.3%,<br>Prefer not to say: 6.8% | 25% | 60% | 60% |
| 2024 | 130+ | 38 | North America: 63%,<br>Asia: 19%,<br>Africa: 12.7%,<br>South America: 3.6% | Asian: 29.5%, White: 28%,<br>Hispanic/Latinx: 24%,<br>Black/African American: 15%,<br>Prefer not to say: 3.5% | 23% | 45% | 51% |

#### Six lectures

get the students up to speed on the engineering design process. They explore the entire creative process, from uncovering customer needs through evidence-based discovery to transforming those insights into actionable design specifications. Students learn how to develop and refine concepts, evaluate ideas using tools like the engineering design decision matrix, and bring their vision to life through prototyping, all while integrating business development strategies to ensure practical and market-ready solutions appropriate for the culture in which their stakeholders reside. These lectures are provided in real-time interactive video presentations as this format has been shown to be more effective than pre-recorded videos for learning new factual information for young adults (De Felice, Vigliocco et al. 2021).

#### Faculty presentations

(called Coffee and Conversations) provide the students with a taste of current research and development in the biomedical engineering field, and they are kept short (~20 minutes) to provide ample time for questions and discussion. The majority of the discussions are led by faculty from our institution, with at least two faculty members per session and each summer at least one is a panel discussion of prominent leaders in the field from other institutions. **Professional development workshops** include biomedical ethics, healthcare disparities, effective communication, reproducibility and rigor, and a career panel. These workshops are led by faculty from both within BME and from other departments within our university.

#### The virtual design projects

topics are related to global health, healthcare in low-resourced environments, and minimizing healthcare disparities. Four-person teams are created based on student interests and availability (due to jobs, time zones, etc.). They are provided with ~10 potential topics and invited to submit their top three choices. In addition to the choices, students indicated the time slots that they are available across different time zones. Based on their interest, weekly availability across time zones, participants are assigned one of those three choices. This approach allowed teams to include participants from multiple countries and continents, maintaining the intended global exposure while accommodating practical scheduling constraints. For example, in 2022, all 30 groups had international students from at least two countries; 23 groups had at least one student from the US. The groups worked together (meet twice a week via online communication tools such as Zoom) through the EDP including web research, primary literature, ideation, materials selection (appropriate for region), and CAD drawings. At the half-way point, each team conducts a critical design review for a BME faculty member to obtain personalized feedback and mentoring. Typically, six BME faculty members are available for live review sessions, during which they meet with student teams and provide targeted feedback. This ensures that every team that requested a meeting received meaningful input, though it required careful scheduling and advance coordination.

#### The final deliverable

for the program is a 4-minute recorded slide presentation from each team which is uploaded to a YouTube “channel” specific for that year. All presentations are reviewed by the organizers and awards for best overall design, best presentation, most creative, and best reduction to practice are given at a final closing ceremony that also includes viewing portions of exemplary presentations and student testimonials. The participants are also asked to review other teams’ presentations and provide “likes” (as appropriate), and the teams with the most likes also earn awards.

#### The time commitment

for the participants is about five hours per week, with three scheduled one-hour components (a lecture, a professional development workshop, and a faculty discussion), and a couple hours of individual and team-based project work. The three scheduled one-hour components are synchronous and interactive to build connections, but each is also recorded and posted on our LMS (Canvas) for students who cannot make a particular time. Certificates of completion are provided for students who attend (or watch asynchronously) 80% of the lectures and discussions and contribute to their team’s design and video presentation. Weekly attendance is monitored on Canvas LMS.

### Reflection

Short-term study abroad programs, as well as international and inter-institutional class collaborations and competitions, offer valuable opportunities for students to step outside their comfort zones, engage in problem-solving within diverse teams, and gain exposure to globalization (Marutschke, Kryssanov et al. 2019). In our program, the students’ ability to work in teams, although separated by vast distances, was evident in the presentations in which the distribution of time between members was generally even, and this was reinforced by peer feedback provided at the end of the program. They also showed impressive ingenuity and cultural sensitivity in their designs. They researched the local cultures (some of the team members coming from those communities) and available resources. One stand-out project involved the use of corn husk to create umbilical cord ties (***Figure 1***, right), with these materials being plentiful and low-cost in the intended market. These findings reinforce previous research on alternative international experiences that demonstrate comparable benefits to traditional study abroad in enhancing participants’global competence while often being more cost-effective and requiring less or no time spent abroad (Yates, Wentz et al. 2020).

**Figure 1:**
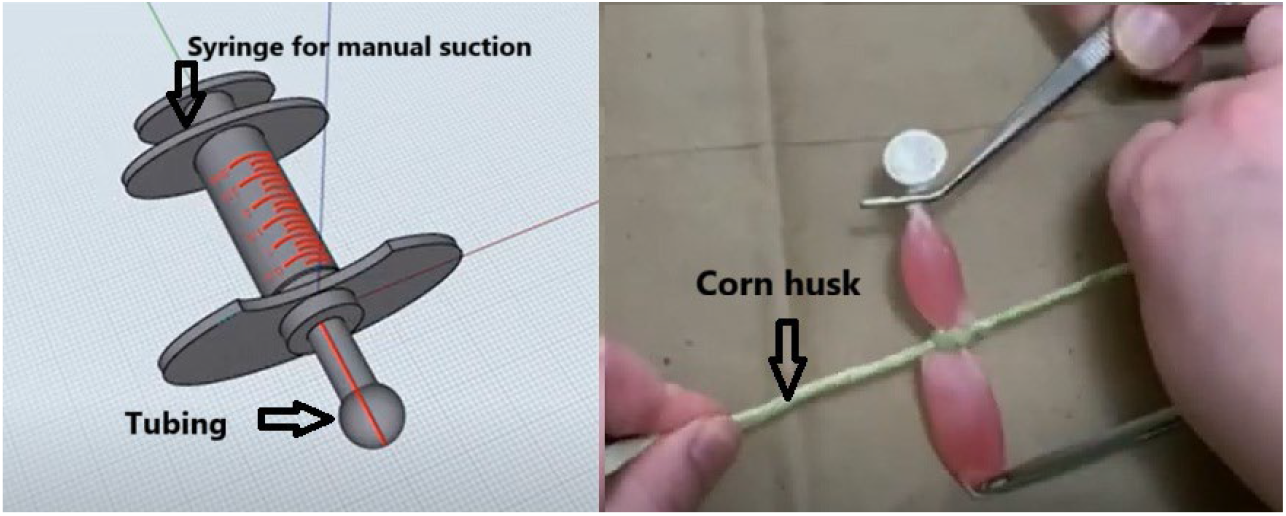
Example pictures from projects that exemplify culturally sensitive design. One team (left) designed a glucose sensing pacifier for painless measurements in babies, and another team (right) designed low-cost umbilical ties made from locally sourced and plentiful corn husk.

Whereas projects in engineering education have traditionally been used as capstones to demonstrate mastery of knowledge and skills, PjBL clearly demonstrates value at an introductory level (Boudreau and Wobbe 2019). Similar to first-year university programs, our program assumes no advanced technical knowledge, yet due to this constructivist approach to learning, students gain valuable problem-solving, interpersonal, and communication skills and demonstrate higher-order thinking through extensive collaboration, iterative problem-solving, and presentation of learning.

Virtual internships, which share many similarities to the virtual design experiences offered to our students, enable large-scale participation in complex problem-solving while still delivering personalized mentored learning experiences. We used the term “virtual internship” in the 1^st^ year of the program, but we found that this terminology was problematic for international students due to specific rules around internships, thus we changed it to “virtual immersion.” Research has shown that virtual internships enhance interest in engineering, particularly among women, and foster the development of engineering design thinking (Chesler, Ruis et al. 2015). Virtual internships are especially advantageous for non-traditional students balancing work and family commitments, as they provide flexibility and accessibility (Ruggiero and Boehm 2016). The COVID-19 pandemic accelerated their adoption, demonstrating their potential to ensure equitable access to experiential learning opportunities even when not able to attend in person regardless of the personal reason (Ekstedt, Ekstedt et al. 2021). Table I shows the strong need for the program, with over 800 students successfully completing the program over four years.

Most students exhibited strong dedication to the program with insightful questions during the Coffee and Conversations, rich discussion during feedback sessions, and having less than 20% of those who signed up not completing the requirements. Further, a search of “VIBE internship” yields hundreds of hits on LinkedIn, demonstrating that students find the experience important enough to put on their resumes and online presence.

### Program Evaluation

#### Qualitative Assessment

The students’ application of the EDP and demonstration of higher-order thinking (i.e., applying knowledge, analyzing alternatives, and creating a solution (Anderson, Krathwohl et al. 2001)) were evident from their impressive presentations at the end of each program. Teams created detailed computer aided designs (CAD), e.g., ***Figure 1 (left)***, and some even created physical prototypes with 3D printing, Arduino microcontrollers, or other materials (***Figure 1, right)***. With the constraints of the virtual environment, and the various student backgrounds (many with only one year of engineering completed), we focused on the engineering design process tools and creative ideation/conceptualization. As students did not, in general, have access to engineering equipment and the skills to use such equipment, this experience is not a substitute for technical design e.g., circuits, mechanical testing, materials processing, etc. Although most teams utilized CAD software to create drawings and some teams were able to create and test physical prototypes, this experience is also not a substitute for senior capstone design.

#### Quantitative Assessment

To more quantitatively assess if our program is meeting its goals for continuous improvement, we analyzed the video transcripts from 52 teams from 2023 and 2024 using NotebookLM (Google, v1.13.0 accessed June-October 2025); these years were analyzed to complement the participant survey data from those years. The YouTube videos were uploaded along with a basic EDP rubric appropriate for an introductory design course which does not include criteria for multiple designs, physical prototypes, or validation and verification (***Supplementary materials***). The LLM was prompted to “evaluate each of the videos using the uploaded rubric and put all of the results into one table with a column for each team’s scores.” Before running the LLM analysis on all of the reports, the authors validated the automated analysis by scoring 10% of the reports (n=5) randomly chosen and comparing the detailed evidence cited for each score to that provided by the LLM. As the scores and the text evidence were very consistent between the authors and the LLM, we proceeded to run the automated analysis multiple times. Although the specific scores varied in a few cases (e.g., a score of 2 in one run and 3 in another for a specific outcome), the average scores were within two points between runs. Our finding of agreement between the LLM and the human evaluator are in line with studies that show that show the potential of LLMs to identify student errors with high accuracy (Bewersdorff, Seßler et al. 2023) and to enhance the accuracy, fairness, and efficiency of grading practices (Fagbohun, Iduwe et al. 2024).

However, our findings are in contrast to a study using LLMs to grade construction management capstone projects in which the LLM generally scored higher than humans, and the overall correlations with two human evaluators were poor, especially with a detailed rubric provided. This discrepancy could be due to complexity and creativity required for the capstone design and the subjectivity in grading “argumentative assignments” (Castelblanco, Cruz-Castro et al. 2024). In terms of data privacy in the AI analysis, NotebookLM does not train its AI models using the uploaded information, and regardless, all videos are publicly available on YouTube.

The automated analysis showed that teams performed well with an average of 87% ±8% (min 71%, max 100%, n=52), all from just a 4-minute video, demonstrating a strong ability to communicate their application of the EDP to solve their chosen problem. The highest scores were earned for Problem Definition & Needs Analysis and Communication (almost all teams 3 out of 3), and the lowest scores with the basic rubric were for concept selection and prototyping (1.3-1.7 out of 3) as they did not have time to discuss multiple concepts in the short format videos and many prototypes were limited to CAD drawings.

#### Student Perceptions

For formative feedback to improve the program, we gathered qualitative feedback from the VIBE participants at the end of the 2023 and 2024 sessions. The [redacted university] Institutional Review Board (IRB) determined that this research is exempt from further IRB review under 45 CFR § 46.104. Participants in 2023 praised the program’s global collaboration, flexible virtual format, and Coffee & Conversations sessions (92% recommendation rate), but criticized Professional Development (PD) sessions for being exclusionary to remote attendees (avg. rating: 3.2/5). Other points included time zone challenges and lack of structured deadlines, with suggestions to improve PD engagement and provide faster access to recordings. Despite these issues, 88% reported a better understanding of low-resource community challenges, highlighting the program’s success in increasing students’ awareness of the importance of taking into account local stakeholder needs when involved in real-world problem solving.

Based on the feedback in 2023 we restructured the PD sessions. In 2024, PD sessions improved slightly (avg. rating: 3.8/5, +0.6), but uneven workload distribution emerged as a new concern, with participants noting a “last-minute crunch.” While global teamwork remained a strength (88% positive, up 3%), group dynamics worsened, with more reports of unresponsive teammates. The 2024 cohort echoed 2023’s call for better accountability tools (e.g., shared task boards) and time-zone-aware scheduling, suggesting these are systemic issues. Despite a minor dip in retention intent (82% vs. 85%), the program maintained high relevance for BME career exploration, with 89% still recommending it.

Student responses to open-ended questions provide a glimpse of the impact of the program. Students highlighted meaningful learning from the program, such as finding value in *“learning about the design process and completing my first design project*.*”* They also noted the value of exposure to different perspectives, saying *“working with students from other universities… allowed for new perspectives and ideas to be created*.*”* The professional development component was another key takeaway, with one student sharing that *“the coffee and conversations provided great insight about potential career choices*.*”* Finally, the program helped them understand how to adapt and succeed in real-world conditions, as one reflected, *“Despite being in a low internet area, I was able to complete necessary work due to the platforms and deadlines given*.*”*

### Challenges

Engaging hundreds of students across multiple institutions, time zones, and languages, presented logistical challenges, including workload equity within design teams. While participation in the group project was required for certification, peer feedback was not included in this iteration; future programs may incorporate it to enhance team accountability. Another aspect that was not explicitly addressed was that while students explored the cultural contexts of their projects through research and discussions, explicit cultural sensitivity training was not provided and represents an opportunity for enhancement in future iterations of the program.

A key consideration for scaling the program was faculty coordination. Scheduling sufficient coverage for design reviews, Coffee & Conversations, and professional development sessions proved challenging, particularly during the summer months when faculty travel was common. Early planning was essential to align availability with program needs, and sustaining consistent engagement across multiple program elements required deliberate effort and recognition of faculty contributions. These factors emerged as the primary choking points for scaling the program. Another aspect that was not explicitly addressed was that while students explored the cultural contexts of their projects through research and discussions, explicit cultural sensitivity training was not provided and represents an opportunity for enhancement in future iterations of the program.

Finally, running a program for so many students takes a substantial effort even if it is scalable and virtual; with the enrollments over the past few years, the program costs $50-$100 per participant as it is critical to pay the instructors equitably. To raise the funds, we obtained mini grants from our institution and a few generous donors stepped up; we are now attempting a crowdfunding effort. Further, as the program scales, more students attend the professional development workshops and can overwhelm the facilitators. After the first year, we found that we needed to create our own workshops that were optimized for virtual attendance and asynchronous viewing.

## Authors’ contributions

KB conceived the program and drafted the manuscript. TA and OA implemented the program in various iterations and edited the manuscript. SM taught the participants engineering design and edited the manuscript.

## Acknowledgements

We acknowledge Prof. Robert Kirsch and Steve Fening of Case Western Reserve University who provided advice to the authors for the VIBE program based on their free virtual summer enrichment program that enrolled ~500 BME undergraduate students in 2020 when almost all summer internships were cancelled due to the COVID19 pandemic. We also acknowledge partial funding for the program from the WPI BME Department, a WPI Summer Sandbox Grant, Edward Mackey, and an anonymous donor.

## Supplemental materials

1. Engineering Design Rubric used to analyze VIBE presentation transcripts
2. Notes about Google NotebookLM analysis with evidence-backed scoring

