## supplementary information for "Virtual Immersion in Biomedical Engineering (VIBE): Exposing undergraduates to culturally sensitive engineering design and professional experiences at scale"

### Supplemental Information:

#### Engineering Design Rubric used to analyze VIBE presentation transcripts.

| Category | 3 | 2 | 1 |
|---|---|---|---|
| **1. Problem Definition & Needs Analysis** | Clear, concise problem statement. Thorough research and stakeholder analysis. Clear constraints and criteria. | Problem defined, some stakeholder input. Constraints and criteria mostly clear. | Vague or incomplete problem definition. Little research or unclear needs. |
| **2. Background Research** | Extensive and relevant research. Sources are well-documented. Incorporates technical and contextual knowledge. | Adequate research, but lacks depth or documentation. | Limited or poorly integrated research. Weak justification for decisions. |
| **3. Design Specifications** | Qualitative and quantitative specifications | Qualitative specifications | Specifications absent or vague. |
| **4. Concept Selection** | Clear and justified criteria used to evaluate options. Logical, data-supported selection. | Criteria used, but selection may be somewhat subjective or weakly justified. | Poorly justified or arbitrary selection. Criteria unclear. |
| **5. Design Development** | Detailed drawings, models, and/or simulations. | Functional design, but lacking depth. | Incomplete or unfeasible design. |
| **6. Prototyping** | Prototype created and described effectively. Virtual is fine. | Prototype created with limited documentation. | Prototype absent |
| **7. Communication** | Design and decisions clearly communicated | Some clarity in communication; | Poorly organized, unclear, or unprofessional communication. |

### Notes about Google NotebookLM analysis with evidence-backed scoring:

When the analysis was run with NotebookLM multiple times, the specific scores varied in a few cases (e.g., a score of 2 in one run and 3 in another for a specific outcome); however, the average scores were within two points between runs. Further, the scores and the text evidence were very consistent between our manual analysis and the LLM. As an example of the evidence cited by NotebookLM for the scoring, for Team 7A from 2023 consisting of four female students from the University of Florida (USA), Cornell University (USA), the Universidad de los Andes (Columbia), U Calgary (Canada) designing a lead filtering device for drinking water, the LLM analysis provided the following evidence for a score of 3 for Criteria 1 (Problem Definition & Needs Analysis): "The video identifies lead poisoning in rural Colombia due to illegal mining as the core problem. It provides a clear needs statement for "indigenous communities in Colombia". Constraints and criteria are defined quantitatively (capacity of at least 40 gallons, sensor level of 10 micrograms per liter) and qualitatively (must filter particulate/dissolved lead, viruses, and bacteria; must be easy and economically efficient to install/maintain)." This criterion was also scored a 3 by the authors for the same reasons, and the LLM scored it a 3 on all three repeated analyses. The total scores for the three repeats were 17, 18, and 17 (out of 21), with one-point differences in scoring between the repeats for three of the criteria: Concept generation, Concept selection, and Design development.
